# Effects of land use and weather on the presence and abundance of mosquito-borne disease vectors in a urban and agricultural landscape in Eastern Ontario, Canada

**DOI:** 10.1101/2021.12.26.474201

**Authors:** Miarisoa Rindra Rakotoarinia, F. Guillaume Blanchet, Dominique Gravel, David Lapen, Patrick A. Leighton, Nicholas H. Ogden, Antoinette Ludwig

## Abstract

Weather and land use can significantly impact mosquito abundance and presence, and by consequence, mosquito-borne disease (MBD) dynamics. Knowledge of vector ecology and mosquito species response to these drivers will help us better predict risk from MBD. In this study, we evaluated and compared the independent and combined effects of weather and land use on mosquito species occurrence and abundance in Eastern Ontario, Canada. Data on occurrence and abundance (245,591 individuals) of 30 mosquito species were obtained from mosquito capture at 85 field sites in 2017 and 2018. Environmental variables were extracted from weather and land use datasets in a 1-km buffer around trapping sites. The relative importance of weather and land use on mosquito abundance (for common species) or occurrence (for all species) was evaluated using multivariate hierarchical statistical models. Models incorporating both weather and land use performed better than models that include weather only for approximately half of species (59% for occurrence model and 50% for abundance model). Mosquito occurrence was mainly associated with temperature whereas abundance was associated with precipitation and temperature combined. Land use was more often associated with abundance than occurrence. For most species, occurrence and abundance were positively associated with forest cover but for some there was a negative association. Occurrence and abundance of some species (47% for occurrence model and 88% for abundance model) were positively associated with wetlands, but negatively associated with urban (*Culiseta melanura* and *Anopheles walkeri*) and agriculture (*An. quadrimaculatus, Cs. minnesotae* and *An. walkeri*) environments. This study provides predictive relationships between weather, land use and mosquito occurrence and abundance for a wide range of species including those that are currently uncommon, yet known as arboviruses vectors. Elucidation of these relationships has the potential to contribute to better prediction of MBD risk, and thus more efficiently targeted prevention and control measures.

## Introduction

Mosquitos have been considered as the most dangerous creatures worldwide [1]. With their widespread occurrence and ability to transmit diseases, they are responsible for hundreds of thousands of deaths each year. Over the past 40 years, in Canada, mosquito populations have evolved in terms of composition, abundance and geographic distribution. There were 74 mosquito species identified as occurring in Canada in an exhaustive list compiled in the 1980s [2]. Recently, six new endemic species have been added to the list: *Ochlerotatus ventrovittis*; *Oc. japonicus*; *Culex salinarius*; *Cx. erraticus*; *Anopheles perplexens*, and *An. crucians* [3, 4]. Furthermore, new exotic invasive species from the United States have been repeatedly detected in southern Ontario. *Aedes aegypti* were detected in 2016 and 2017 in southern Ontario. These are though to be repeated introductions, with no evidence of over-winter survival, however the more cold-tolerant *Ae. albopictus* is now thought to have become established [5].

Arboviruses endemic to Canada include West Nile Virus (WNV), Eastern Equine Encephalitis virus (EEEV) and Californian serogroup viruses (CSV) belonging to the Bunyaviridae, including Jamestown Canyon virus (JCV) and Snowshoe Hare virus (SSHV). Three mosquito species are mainly responsible for the transmission of WNV in eastern Canada: *Aedes vexans* and the two species of the *Culex pipiens* and *Cx. restuans*, which are often enumerated together due to difficulty in distinguishing them [6]. *Culiseta melanura* is the main vector for EEEV [7, 8]. CSV have several potential vectors including *Ochlerotatus* and *Culiseta* spp. [9–21]. Other mosquito species present in Canada may also be involved in the transmission of arboviruses as suggested by laboratory studies [22–24]. Therefore, monitoring communities of mosquito species, rather than single species, may be more effective for understanding the ecology of arboviruses and predicting risk from them.

Many studies have highlighted the importance of weather and climate as determinants of the spatio-temporal distribution and abundance of mosquitoes [25–28]. Climate change [29], may be driving current expansions in the geographical range of mosquitoes, particularly northern expansion of ranges [30–32]. Weather and climate impact the occurrence and abundance of mosquitos in multiple ways [33]. Increasing temperature can lead to faster inter-stadial development and shorter life cycles. However, the association between weather and mosquito occurrence as well as abundance is not straightforward. High temperatures above 35 °C kills most mosquitoes species [34, 35] but higher average temperatures could also increase the winter survival rate of eggs and promote earlier hatching, and prolong the season of mosquito activity [36]. Increased rainfall may increase reproduction and abundance by increasing suitable larval habitats. Conversely, excessive rainfall can also decrease abundance via egg destruction and leaching of larvae [37].

Weather and climate are not the sole factors influencing species distributions, abundance and diversity. Land use change related to agriculture and urban expansion, and increased forest loss is the largest driver of landscape modification worldwide [38, 39]. Several studies have linked land cover and land use change to mosquito community changes [40–49]. Urban greening, as a result of urban conservation initiatives, will likely enhance resources for arthropods including mosquitoes [50, 51]. Increased crop irrigation in agricultural environments and associated drainage networks may increase larval habitat for several mosquito species. Habitat structure determines mosquito breeding and resting sites, and nutrition of the different mosquito developmental stages [49, 50, 52–54]. Thus, attempting to understand how land use activities, as well as climate and weather, influence composition and abundance of mosquitos is essential to understand the ecology of mosquito species.

Most studies on mosquitos [25, 55–58] or MBDs [59–63] focus either on weather and/or climate and only occasionally account for land use [40, 64, 65]. Despite evidence that integrated study of weather and/or climate and land use effects can help us better understand mosquito occurrence, abundance and range expansion [66, 67], few studies have done so [68–71]. Furthermore, studies tend to focus on one or a few species and ignore the rarer ones even though the community of vectors as a whole may be important in MBD dynamics [72]. In this study, we aim first to assess the combined impacts of land use and weather variables as determinants of mosquito occurrence and abundance, and to develop a more community-wide approach to assess the risk represented by mosquito species endemic to regions where MBDs have an important impact on public health. Individual differences in species traits were included in this study as they can influence the species response to the explanatory variables. As a first cut analysis, we used readily available remote sensing and meteorological data. Much finer analyzes may follow using micro-climatic and micro-land use data that should improve models accuracy.

## Materials and methods

### Study region

The study sites are located in the greater Ottawa area and the adjacent South Nation river catchment (SN) in eastern Ontario, Canada (Fig 1). The city of Ottawa covers 2,778.13 km^2^ with ∼1 million inhabitants, while the SN river basin covers an area of ∼3900 km^2^ with an estimated population of approximately 115,000. The SN study area is predominately agricultural land, with patchy forested areas and some suburban locations [73]. The climate is semi-continental with a warm, humid summer and a very cold winter [74].

**Fig 1.**
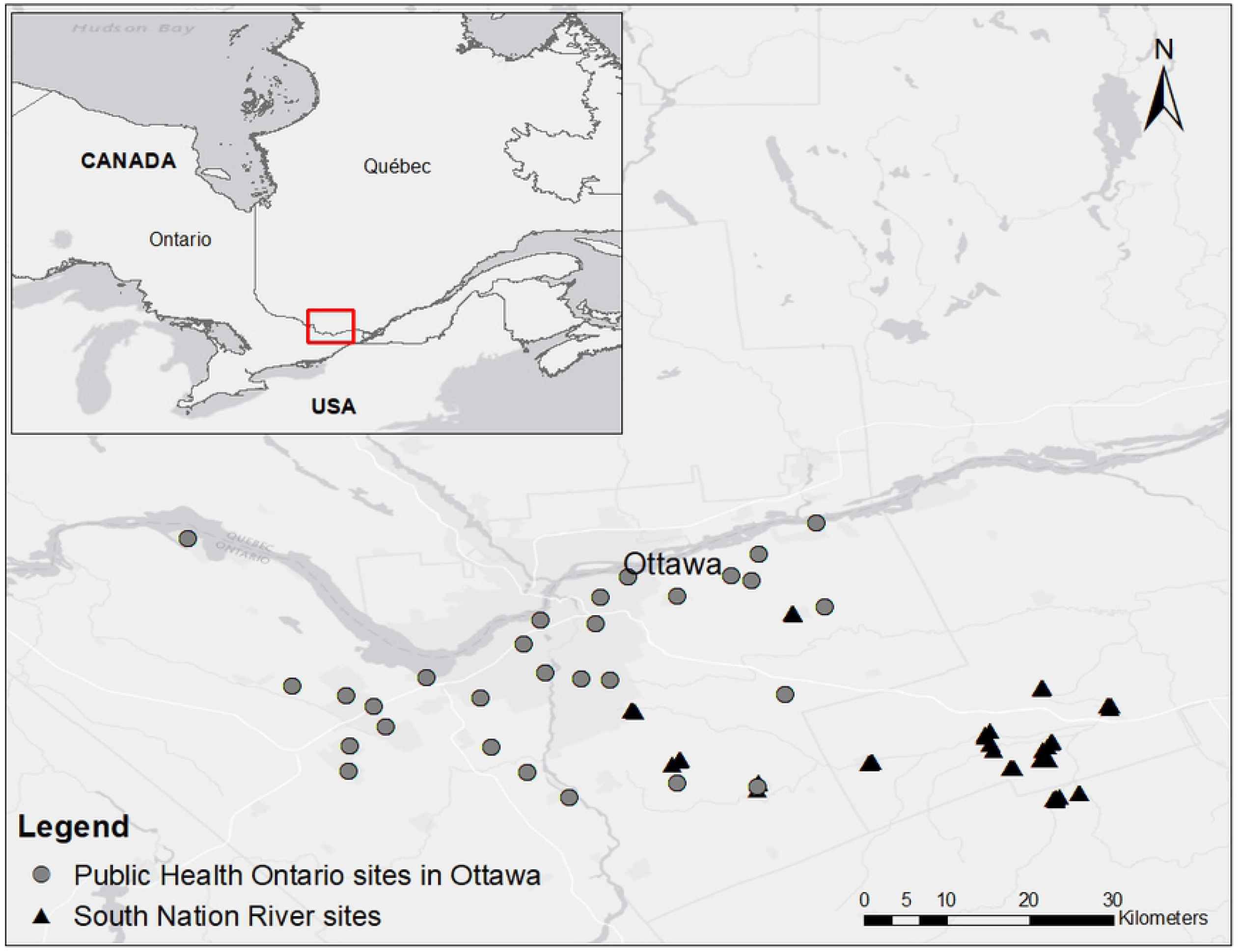
Study region and sampling site locations (Public Health Ontario in Ottawa (29 sites) and the South Nation River watershed (56 sub-sites)).

### Mosquito variables, sampling design and mosquito identification

Mosquito data came from two sources. Prospective field study was undertaken at 56 SN field sites in 2017 and 2018. Data were also obtained from the ongoing West Nile virus surveillance program of the Ottawa Health Unit during 2017 and 2018, in which mosquitoes were captured at 29 sites. At the SN sites, a stratified sampling design was used. Traps were set in locations with (1) suburban and, (2) agricultural land uses class, that each contained seven sites. Each site regrouped four sub-sites according to the presence of water and woodlot features: (1) an area with no water and no wooded area, (2) an area with water but no woods, (3) an area with water that is partially wooded and (4) an area with water that is densely wooded. The number of traps employed by a public health unit in Ontario at a given location varied significantly from year to year. To ensure comparability between the two datasets, we kept only the traps that were in operation during a minimum of 15 weeks. For both datasets, mosquitoes were collected on each site one night every other week from May to end of September (end of August for 2017 in the SN sites), using CO_2_ baited Centers for Disease Control and Prevention (CDC) miniature light-traps. Trapped mosquitoes were identified and counted by species. *Cx. pipiens* and *Cx. restuans* were lumped into a single group *Cx. pipiens-restuans* due to their morphological similarity

### Weather variables

Temperature (minimum and maximum) and precipitation during the study period were extracted from a gridded weather data : Daymet (Daily Surface Weather And Climatological Summaries) (https://daymet.ornl.gov/) [75]. These data are available at a daily time step across a 1km x 1km spatial resolution. Weather data were associated with mosquito trapping locations by averaging weather conditions in a one-kilometer radius around each trapping site. Given that larval development to adult stage can occur between 5 to 90 days [76, 77], these daily variables were averaged over different time period: 5, 30 and 90 days before the capture. With the weather during the day of capture, a total of 12 weather variables were used as explanatory in the models (S3 Table).

### Land use variables

A buffer zone with a radius of 1 km was created around each trapping site, to approximate the average flight range of the 27 studied species [78, 79]. The percent coverage of land use classes was extracted from annual crop inventory data for 2017 and 2018 [78], within each buffer. Crop inventory data were reclassified into seven land use variables (S4 Table).

### Mosquito traits

In this study, we also studied how traits influenced the species response to the environment [80]. The traits, included as a binary variable, were five important behavioral characteristics related to mosquito life cycle and mosquito habitat: 1) number of generations per year, 2) stage of the life cycle that overwinters, 3) type of substrate where the eggs are laid, 4) resistance of the egg to desiccation and 5) typical larval habitat (S6 Table).

### Statistical methods

Where possible we used either presence-absence or abundance of each species detected as response variables. However, for species with low abundance (<10 individuals), only presence-absence models were used. The temporal unit was a period of two weeks (S1 Table), species abundance was averaged biweekly for each trapping site (S7 Fig). The temporal unit as well as the sample identifier were used as random effects to account for repeated sampling from the same site.

Spatial autocorrelation was taken into account using the distance-based Moran’s eigenvector maps (db-MEM) [81]. This method is a spectral decomposition of the spatial coordinates of the sampling locations and allows to account for the proximity of certain sampling locations. The db-MEMs that significantly characterize spatial autocorrelation were included as covariates in the model. Construction and test of db-MEM were performed using the adespatial R package [81].

Relationships between mosquito occurrence or abundance and environmental variables were explored using a Hierarchical Modeling of species Community (HMSC) analysis [80]. Mosquito abundance data were explored using over-dispersed Poisson models, whereas for mosquito presence/absence data probit models were used. HMSC model parameters are estimated using a Gibbs sampler. For the probit model, the MCMC chain was run for 300,000 and 320,000 iterations, for weather-only and weather-and-land-use models respectively, half of which were burn-in iterations. For the over-dispersed Poisson model, weather-only and weather-and-land-use models were run for 200,000 and 220,000 iterations respectively, half of which were burn-in iterations. Convergence was assessed using the Geweke’s convergence diagnostics [82]. Convergence was assumed to be reached for a parameter if it had a Geweke diagnostic value lower than two. Trace and density plots for all parameters were also checked visually to ensure convergence was reached.

Adjusted Efron’s pseudo-R^2^ was computed to compare the occurrence and abundance models. This metric is a relative measure of model fit, obtained by calculating the proportion of explained variance over the total variance in the data [83] adjusted for the number of parameters in the model. We also assessed the predictive performance of occurrence-based models by using the area under the receiver operating characteristic curve (AUC) and the root-mean-squared error (RMSE). AUC evaluates the ability of the model to discriminate accurately occurrence and absence. We used the RMSE and the correlation between predicted and observed abundance for the abundance models. All multivariate models were carried out with the “HMSC” package [84] through the R statistical language version 4.0.2 [85].

Models with and without land use were compared by looking at the values of their respective adjusted pseudo-R^2^. The explained variation was partitioned into components related to fixed effects (weather, land use, spatial component) and random effects (two weeks’ period and sample identification) for each species, following the procedure proposed by Ovaskainen et al. (2017) [80].

## Results

### Mosquito sampling

Thirty species belonging to seven genera (*Aedes (Ae), Anopheles (An), Coquillettidia (Cq), Culiseta (Cs), Culex (Cx), Ochlerotatus (Oc), Psorophora (Ps)*) were detected with a total of 245,591 adult female mosquitos captured. The biweekly average abundance of each species varied over the two years of collection (Fig 2). The total abundance of all combined species was higher in 2017 (163,211) than in 2018 (81,628).

**Fig 2.**
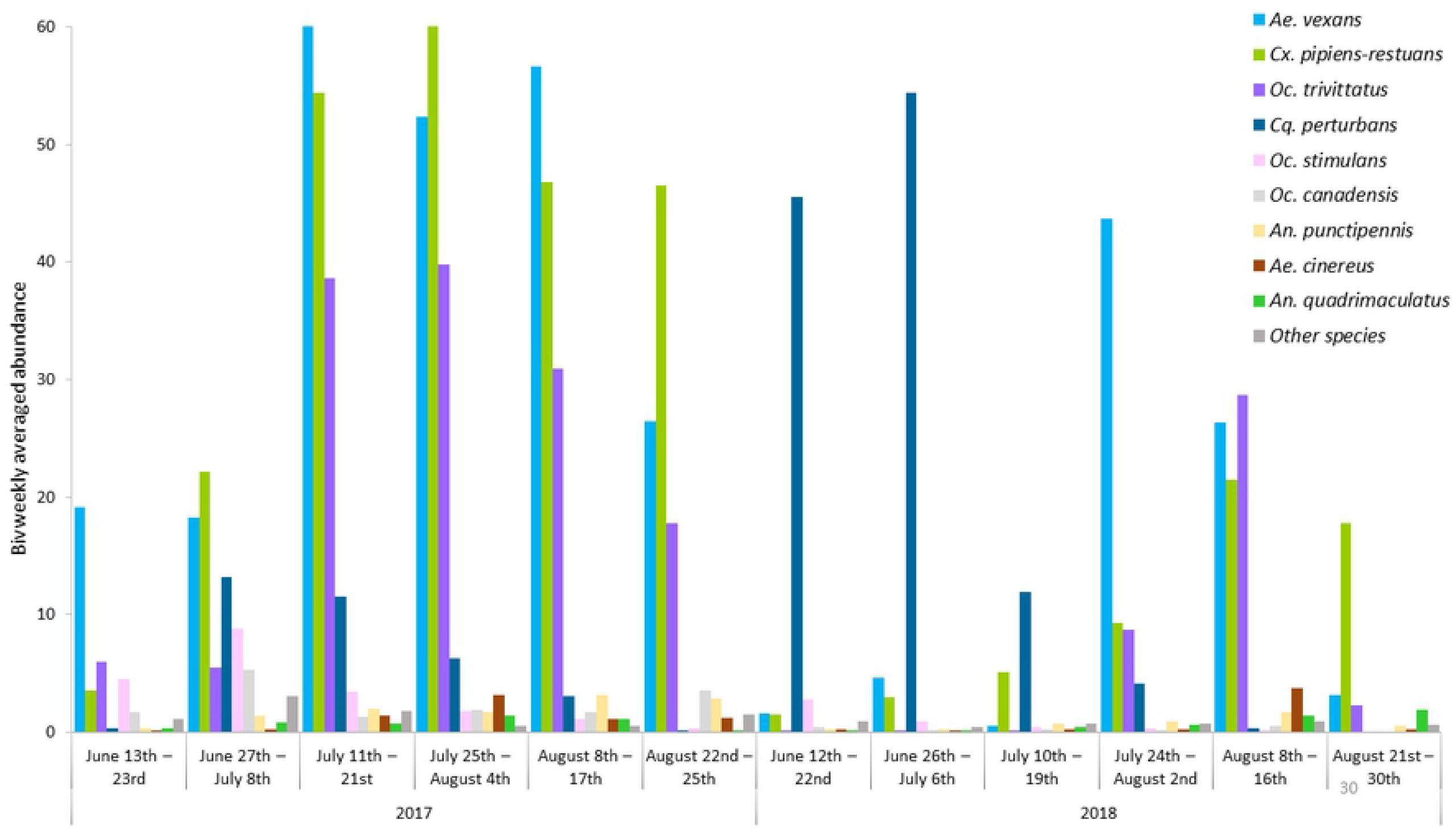
Seasonal changes in biweekly averaged abundance for all species over 2017 and 2018 collection period in Ottawa and the South Nation river sites. The “other species” are rare species with average abundance less than 1. They include *An. barberi, An. walkeri, Cs. melanura, Cs. minnesotae, Cs. morsitatns, Cx. salinarius, Cx. territans, Oc. abserratus, Oc. cantator, Oc. communis, Oc. dorsalis, Oc. excrucians, Oc. fitchii, Oc. intrudens, Oc. japonicus, Oc. provocans, Oc. punctor, Oc. sticticus, Oc. triseriatus, Ps. ciliata* and *Ps. ferox*.

### Added value of land use as a predictor of mosquito occurrence and abundance

Of the thirty species observed, only 27 were included in statistical analysis due to lack of information on the traits of three species (*An. barberi, Oc. sticticus* and *Cx. salinarius*). All models for all species converged. Density plots are provided in the supporting information (S17-20 Fig). The community-level explained variation, illustrated by the mean-values of the species-specific adjusted Efron’s pseudo-R^2^, was computed for each model ranging from 0.21 to 0.28 (Table 1). The values of the community-level adjusted Efron’s pseudo-R^2^ suggest that including land-use in the model in addition to weather can modestly improve overall predictive power. However, because the community-level explained variation is the average R^2^ across all species, it hides the significant variation in independent species (from 0.053 to 0.988 for the occurrence model and from 0.040 to 0.725 for the abundance) (Fig 3). For some species, the adjusted Efron’s pseudo-R^2^ was negative, indicating that these models are less likely than sampling a random variable from a probability distribution, so results from these species are not interpreted.

**Table 1.** Community-level explained variance of the four models.

| | Community-level adjusted pseudo-$R^2$ of weather-only model | Community-level adjusted pseudo-$R^2$ of weather-and-land-use model |
| --- | --- | --- |
| **Occurrence (27 species)** | 0.26 | 0.28 |
| **Abundance (12 species)** | 0.21 | 0.22 |

**Fig 3.**
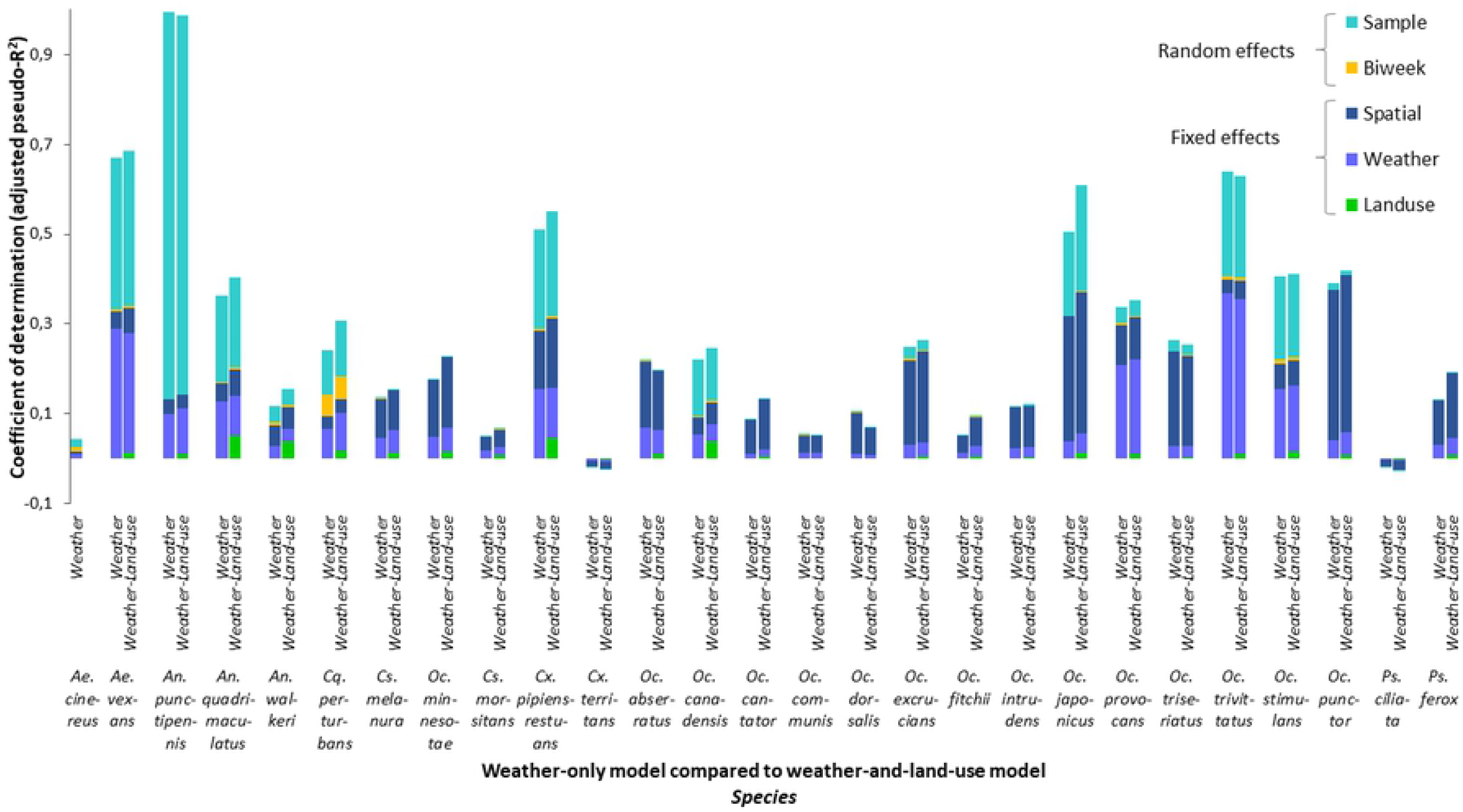
Variation partitioning according to species among weather, land use, spatial fixed effect and the temporal and spatial unit random effects from the occurrence-model. Community-level adjusted Efron’s pseudo-R^2^ of weather-only model (0.24) was compared to that of weather-and-land-use model (0.28). Each bar corresponds to one species.

When variation was partitioned according to each species, there was an increase of the adjusted Efron’s pseudo-R^2^ values in 59% of species from the occurrence model and 50% of species from the abundance model (Fig 3 and 4). This suggests that adding the land use component does not necessarily improve the quality of the models and this is true for both occurrence and abundance. The community-level adjusted pseudo-R^2^ corresponding to land use was low (0.013 and 0.015 for occurrence and abundance, respectively) compared to that of the weather variable (0.069 and 0.057 for occurrence and abundance respectively). In all cases, weather information was most crucial for predicting mosquito occurrence and abundance data in the region.

**Fig 4.**
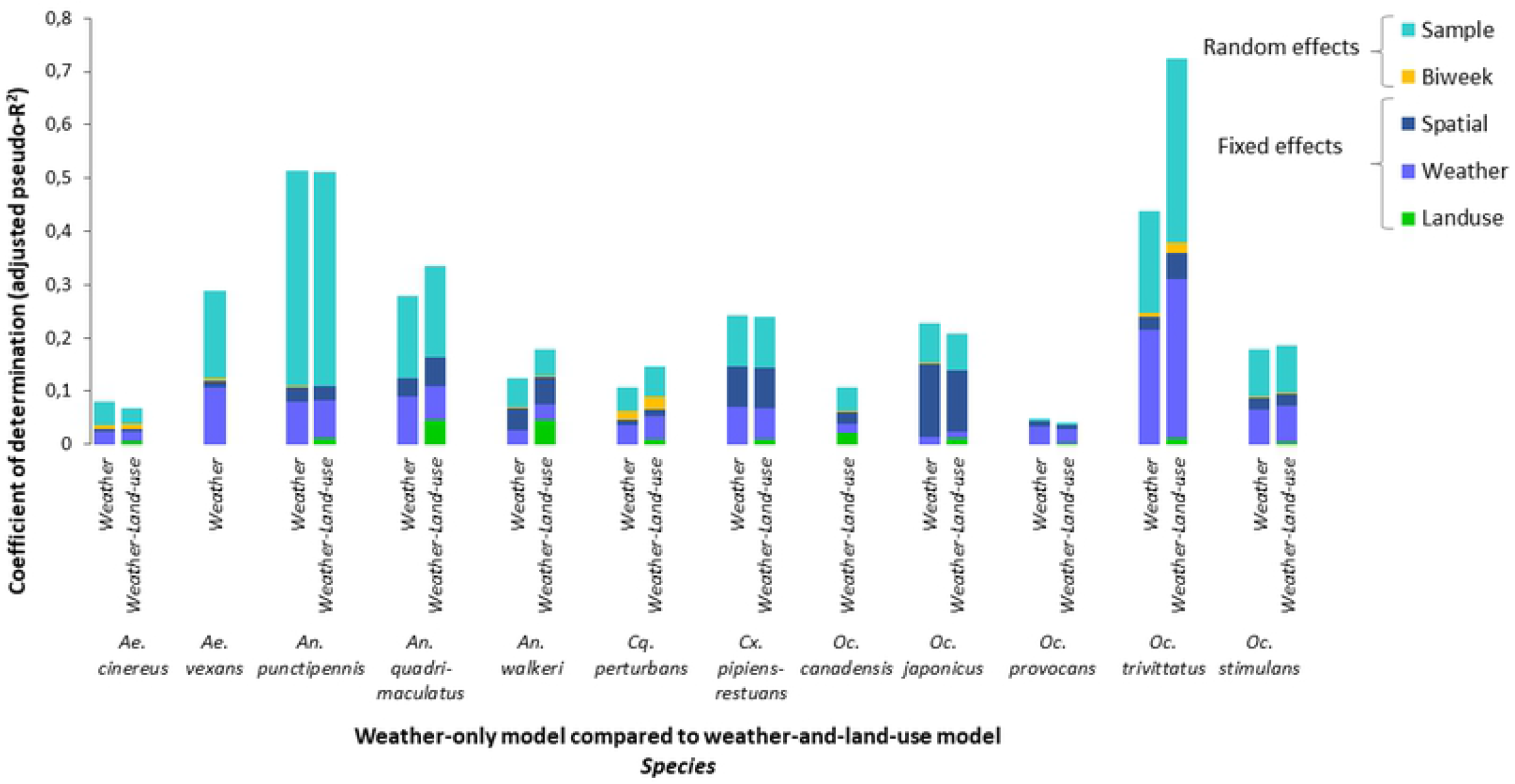
Variation partitioning according to species among weather, land use, spatial fixed effect and the temporal and spatial unit random effects from the abundance-model. Community-level adjusted Efron’s pseudo-R^2^ of weather-only model (0.21) was compared to that of weather-and-land-use model (0.22). Each bar corresponds to one species.

The results also highlight that other components of the models may play important roles in explaining the response variables. For example, approximately half of the whole variation explained was attributed to the sample random effect (10%) suggesting that the sample have relatively strong effects on patterns of species occurrence and abundance while the spatial fixed effect variance ranged between 3.66 and 8.37%. However, the community-level adjusted Efron’s pseudo-R^2^ corresponding to the temporal unit (“Biweek”) remained very small for all species (<1%). The comparison between the occurrence-based model and the abundance-based model revealed that modeling occurrence explains more variation in the data (Community-level adjusted Efron’s pseudo-R^2^ = 0.28) than the models for abundance (Community-level adjusted Efron’s pseudo-R^2^ = 0.22) except for two species (*Oc. trivittatus* and *An. walkeri*). Finally, the proportion of variance related to each group of covariates and random variables was relatively similar for the same species whether in the occurrence model or in the abundance model.

### Weather predictors of mosquito occurrence and abundance

As our primary goal was to evaluate the added value of including land use when modeling mosquito occurrence and abundance, we only present the results obtained from weather-and-land-use models. However, results of the weather-only models are presented in the supporting information (S12 and 13 Fig). Furthermore, since the community-level adjusted Efron’s pseudo-R^2^ does not reflect precisely the variation in the coefficient of determination specific to each species, from here, we focus only on species-level results.

Model results indicated that various weather variables influence occurrence and abundance of a mosquito species (Table 2, S8 and 9 Fig) with weather variables averaged over one to three months prior to mosquito capture being those most important.

**Table 2.**
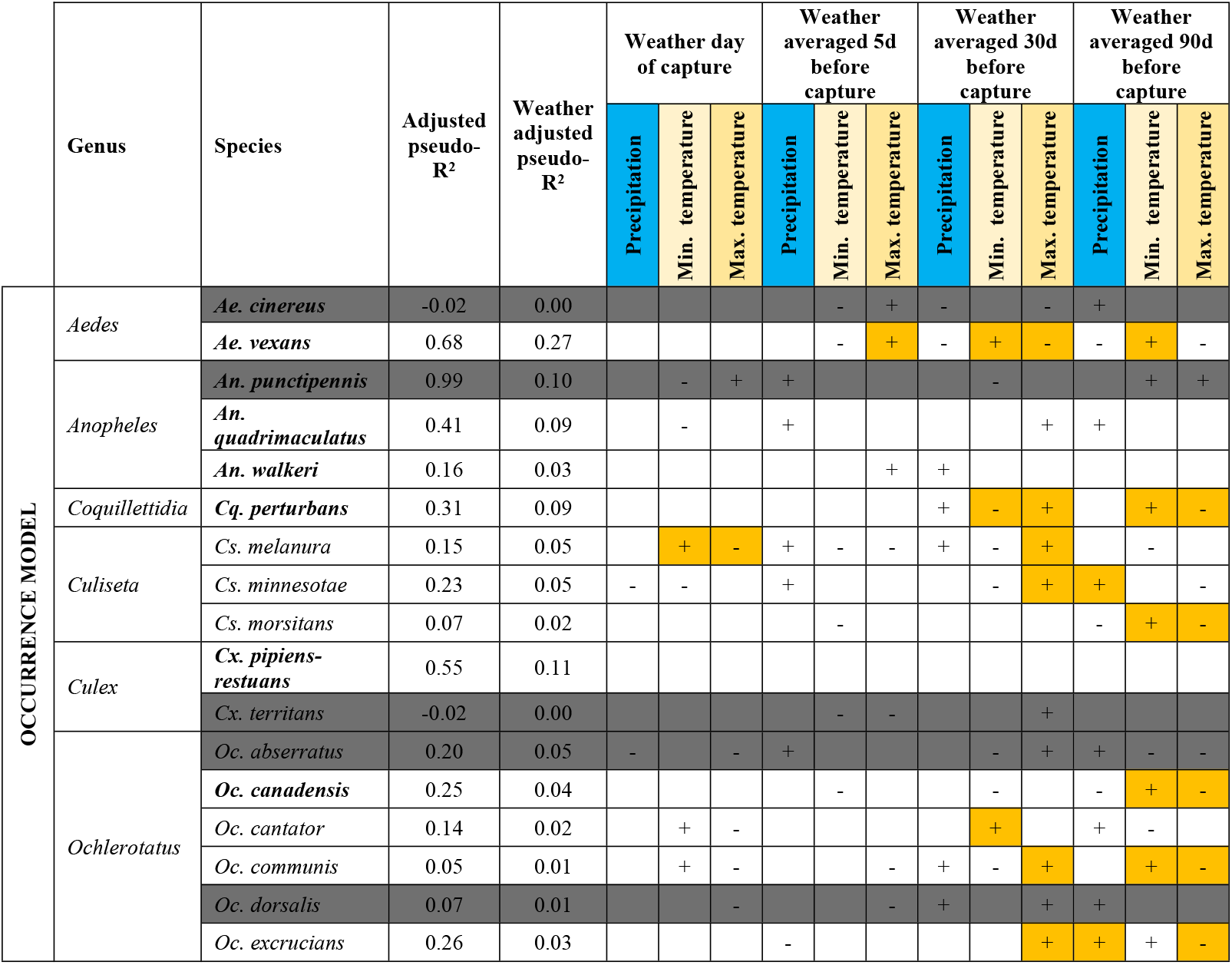
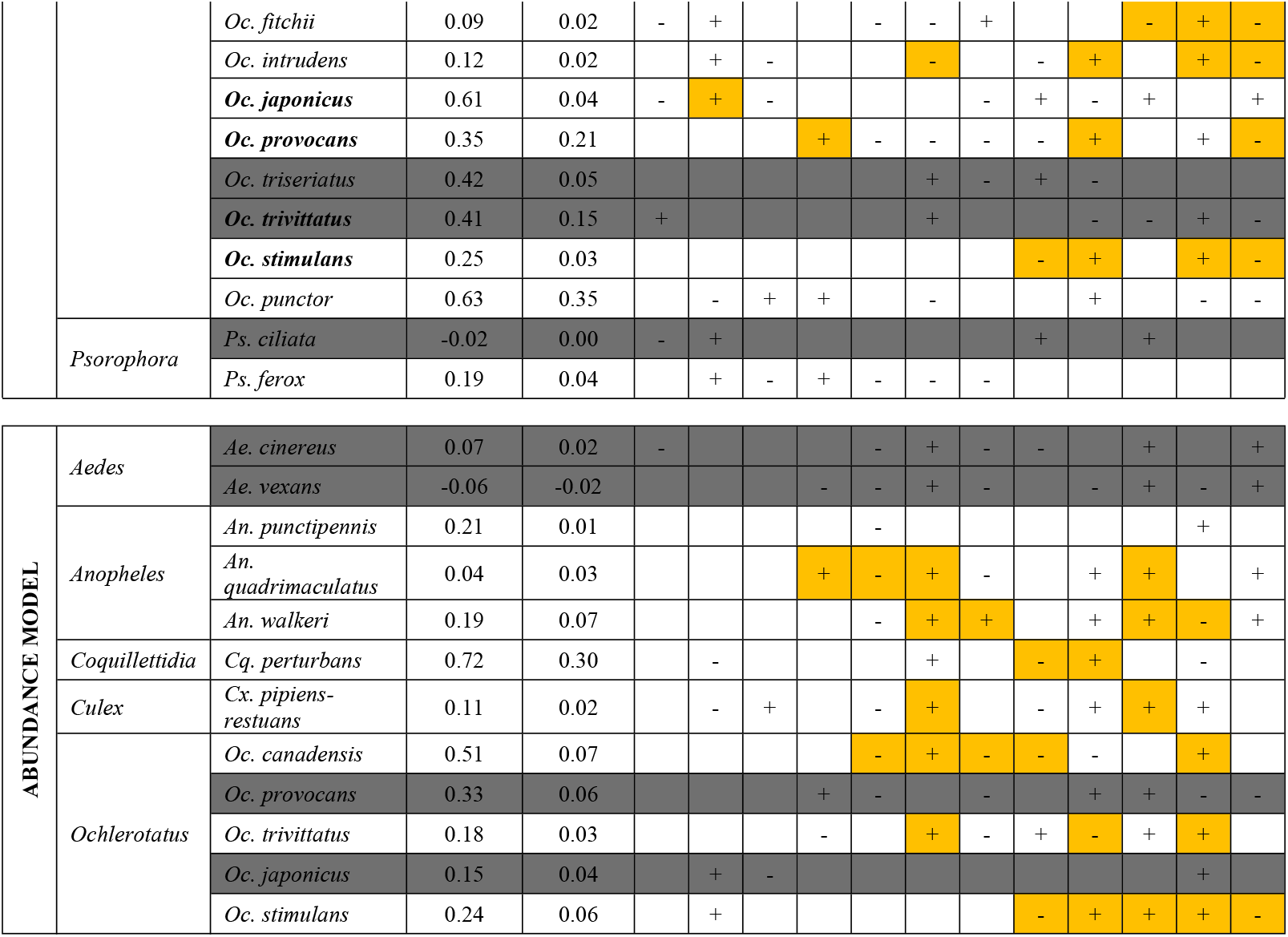
Comparison of the significant and relatively strong effects of weather on mosquito occurrence and abundance of the 12 commonest species. + and - signs represent a positive (average of the marginal distribution of the model parameter above zero) or a negative (average of the marginal distribution of the model parameter below zero) association between the variables and mosquito occurrence or abundance. Orange cells show relationships between weather variables and the response variables (occurrence / abundance) that are most precise (narrow distribution) and important. Grey cells represent species that showed either a negative value of the adjusted Efron’s pseudo-R^2^ or a deterioration of model goodness of fit once land use was added.

#### Precipitation

Occurrence and abundance displayed generally positive responses to precipitation. Precipitation had a greater effect when it was averaged over a large number of days prior to sampling relative to a smaller number of days, especially for the abundance models. Total precipitation during the day of capture did not have a major effect. The occurrence of only *Oc. provocans*, and the abundance of only *An. quadrimaculatus* were positively associated with precipitation averaged over 5 days before capture (Prcp_5_). Precipitation averaged over 30 days had a positive relationship with *An. walkeri* and a negative relationship with *Oc. canadensis*. Precipitation averaged over 90 days was positively associated with the abundance of *An. quadrimaculatus, An. walkeri, Cx. pipiens-restuans* and *Oc. stimulans*, and the occurence of *Cs. minnesotae* and *Oc. excrucians*. The occurrence of *Oc, fitchii* was negatively associated with the precipitation averaged over 90 days.

#### Minimum temperature

The effect of the minimum temperature differed according to the time lag before capture. The minimum temperature averaged over 90 days before capture (Tmin_90_) was a significant determinant for occurrence in 42% of species. The minimum temperature averaged over 30 days (Tmin_30_) was positively associated with the occurrence of *Ae. vexans* and *Oc. cantator* and negatively associated with the occurrence of *Cq. perturbans* and *Oc. stimulans*. In the abundance model, the effect of Tmin_30_ was negative for 38% of species. The minimum temperature averaged over 5 days (Tmin_5_) was a determinant of abundance for only two species (*An. quadrimaculatus* and *Oc. canadensis*) but with a negative effect. Finally, the minimum temperature of the day of capture was positively associated with the occurrence of *Cs. melanura* and *Oc. japonicus*.

#### Maximum temperature

The maximum temperature was significantly associated with mosquito occurrence and abundance. In the occurrence model, the maximum temperature averaged over 30 days (Tmax_30)_ was positively associated with nine of 19 species, but negatively associated with *Ae. Vexans*. In addition, Tmax_30_ was positively associated with the abundance of *Cq. perturbans* and *Oc. stimulans* and negatively associated with the abundance of *Oc. trivittatus*. In the abundance model, the maximum temperature averaged over 5 days (Tmax_5_) stood out as the variable most frequently associated with mosquito abundance with a positive association with five of eight species. The maximum temperature of the day of capture was negatively associated with the occurrence of *Cs. melanura*.

### Land use predictors of mosquito occurrence and abundance

Land use variables had generally weaker associations with mosquito occurrence and abundance than weather variables as shown by the low adjusted Efron’s pseudo-R^2^ values for the land use component. We nevertheless decided to retain the results of the species for which an improvement in the model fit was observed.

Overall, the land use effects were consistent between occurrence and abundance models. The major land use drivers common to all mosquito species were wetland, forest and bare ground (Table 3). There was a positive relationship between wetland coverage and mosquito occurrence (for 9 of 19 species) and abundance (for 7 of 8 species). Coverage of the forest land use class had a positive relationship with mosquito occurrence (for 7 of 19 species) and abundance (for 3 of 8 species) with the exception of *Cx pipiens-restuans* in the occurrence and abundance model and *An. quadrimacultatus, An. walkeri* in the abundance model. Coverage with bare ground was negatively associated with most species (12 of 19 species from the occurrence model and 4 of 8 species for the abundance model).

**Table 3.**
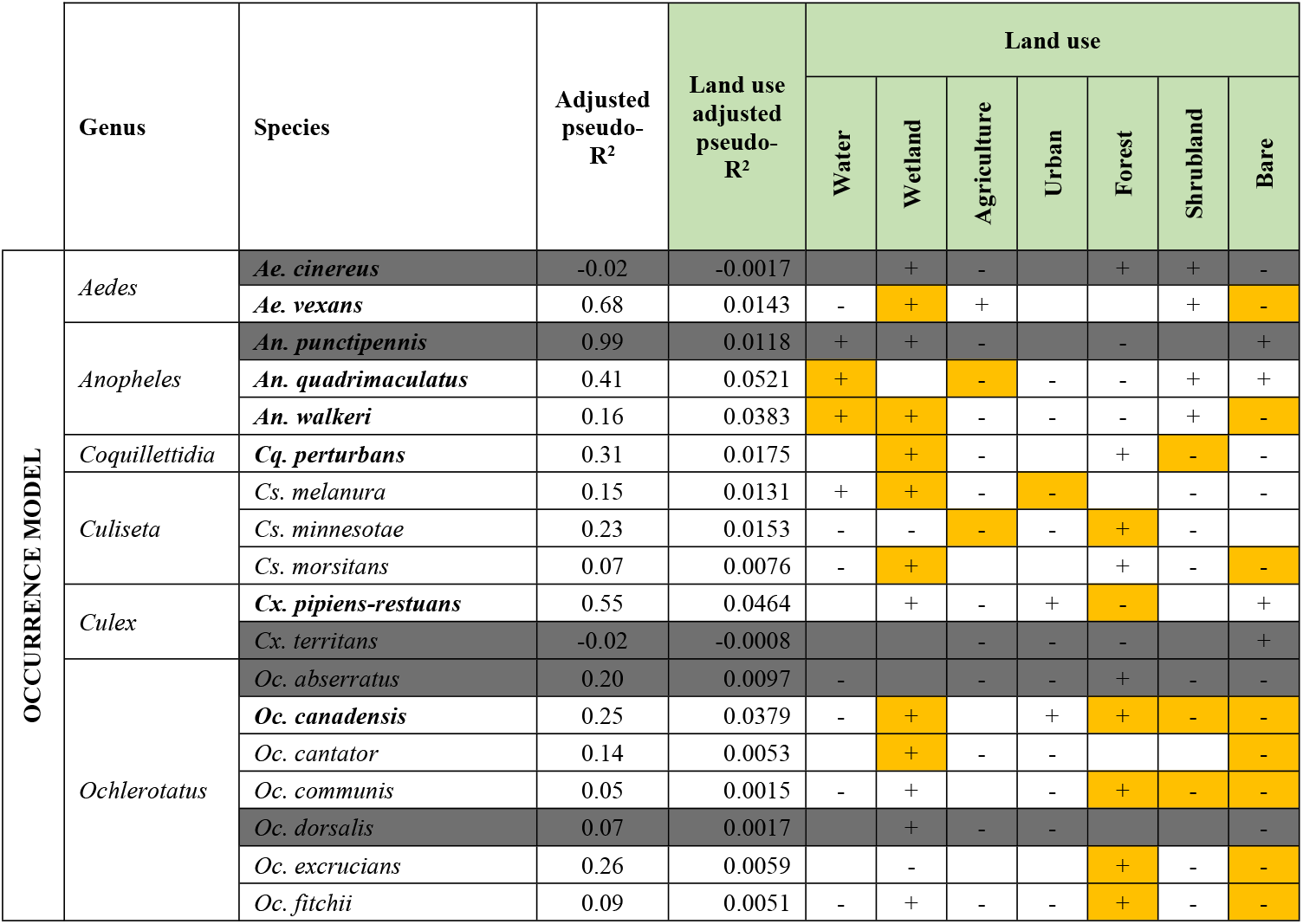
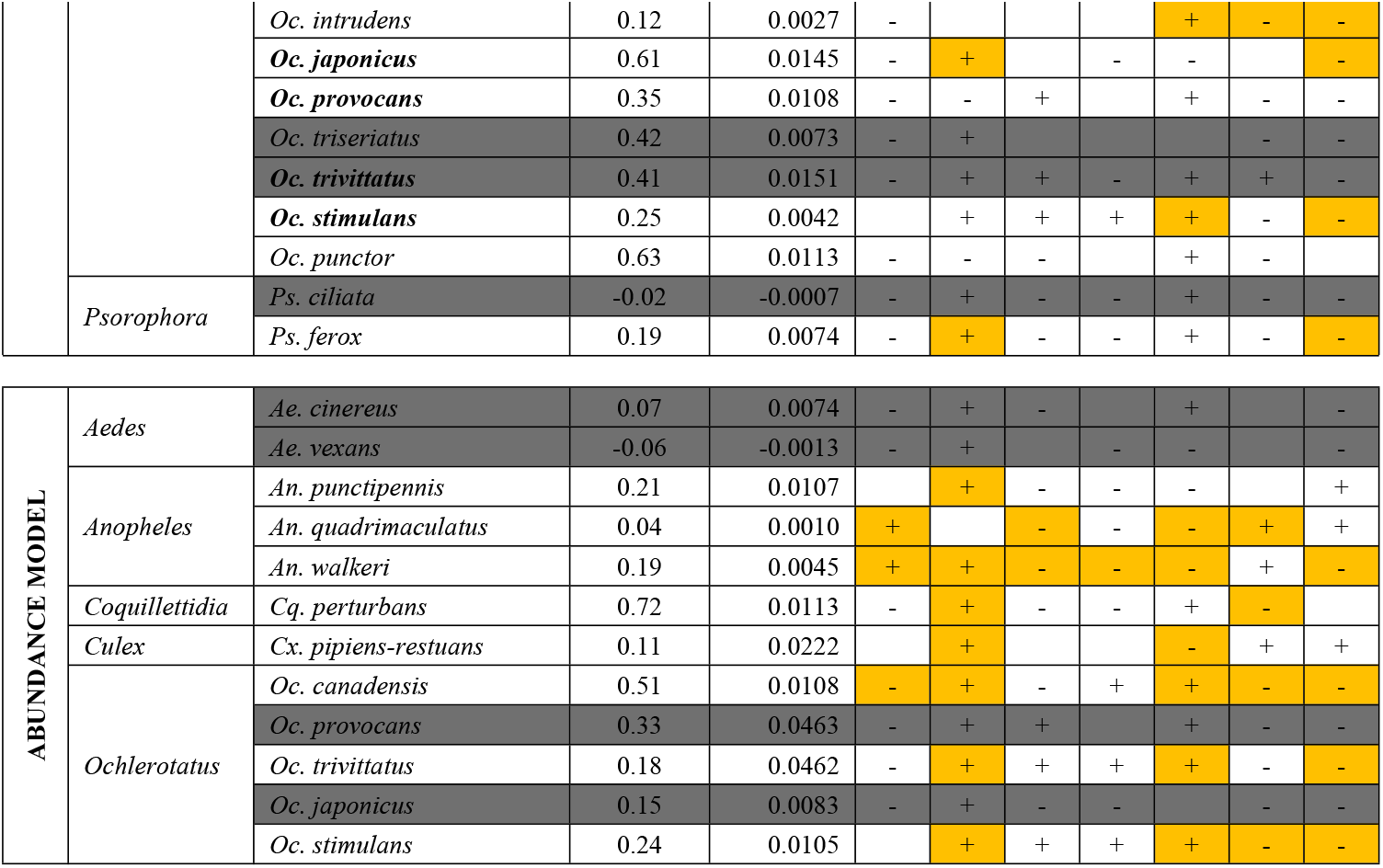
Comparison of the significant and relatively strong effects of land use on mosquito occurrence and abundance of the 12 most common species. + and - signs represent a positive (average of the marginal distribution of the model parameter above zero) or a negative (average of the marginal distribution of the model parameter below zero) association between the variables and the mosquito occurrence or abundance. Orange cells show the relationships between the weather variables and the response variables (occurrence / abundance) that are more precise and important. Grey cells represent species that showed either a negative value of the adjusted Efron’s pseudo-R2 or a deterioration of model goodness of fit once land use was added.

Other land use classes (water, agriculture, urban and shrubland) were infrequently associated with mosquito occurrence and abundance. Coverage with water was positively associated with occurrence and abundance of *Anopheles* species, and negatively associated with occurrence of *Oc. canadensis*. There was a negative relationship between coverage with the agriculture land class and occurrence of *An. quadrimaculatus* and *Cs. minnesotae*, and the abundance of *An. quadrimaculatus* and *An. walkeri*. Coverage with the urban land class had a negative relationship with the occurrence of *Cs. melanura* and the abundance of *An. walkeri*. The shrubland land class was negatively associated with the occurrence of four species (21%) and the abundance of three species (38%) but was positively associated with abundance of *An. quadrimaculatus*.

## Discussion

This study confirms the central role of weather in the ecology of mosquito in Eastern Canada, while highlighting the importance of land use in determining the occurrence and abundance of certain species. By jointly analyzing data from multiple species using the HMSC approach, we were able to gain unique insight into ecological drivers of species that are less abundant and therefore infrequently studied, yet are known vectors for several important mosquito-borne diseases.

### The importance of cumulative weather effects

Our results clearly demonstrate the importance of the accumulation over several days, or weeks to months of favorable climatic conditions for the development of mosquitoes. Mosquitoes are ectothermic organisms, each developmental stage may be sensitive to weather conditions, and there is a species-specific range of temperature conditions that permit completion of their life cycle and, if we are in the optimum of the range, is the shortest possible [86]. However, for univoltine species, i.e. having only one generation per year, high temperature for one month may speed up the life cycle which leads to an earlier emergence of the adults. As adult lifespan can last few weeks at most [77], the cohorts that appeared after one month are no longer present after 90 days. Indeed, we observed that, together with the positive effect of a higher temperature averaged over 30 days before capture, we often saw a negative association when a longer period is considered, i.e. 90 days.

Weather variables that affect mosquito occurrence are not necessarily the same that affect abundance. The importance of precipitation, especially accumulated over several weeks before mosquito capture, appears only in our abundance models. This cumulative effect of precipitation has to be considered alongside the time period between egg laying and adult emergence. The multiplication of mosquitoes depends closely on the availability of breeding sites favored by rainfall as well as the presence of water in these same sites, whether natural or artificial. If the water evaporates quickly, this will give less time for larvae to develop and limit the number of emerging adults [87].

Lastly, warm temperatures on the day of capture had a limited impact on mosquito abundance, compared to the cumulative effects of temperatures over several days. However, for some species, optimal temperatures for adult daily activity apparently remained relatively important, and not in the expected direction. Two species, *Oc. japonicus* and *Cs. melanura*, were positively associated with a warm minimal temperature on the day of the capture. The scarce information from the existing literature indicates that *Cs. melanura* is nocturnal [88] and therefore, adults may be more active during the night when it is generally cooler (Tmin_0_) than during the day (Tmax_0_), and making warm nights potentially favorable to *Cs. melanura*. Explanations for *Ae. japonicus* are less obvious. This species is known to be cold tolerant [89], and may be more active during cool days (intermediate temperatures).

An apparent limitation of our study was to use the weather as a proxy of mosquito seasonality. However, mosquitoes don’t respond instantaneously to weather but accumulate past effects of this latter to develop cohorts over the season. Using the average weather over previous time interval partly solves this issue but our approach doesn’t represent a seasonal model per se. Our model will still predict high abundance of mosquitos in early or late summer where weather conditions are exceptionally favorable, no matter if the population had enough time to develop. Thus, our study highlights the need of a seasonal model for community, currently non-existent, that will be constructed on population dynamics to reflect this cumulative build up of the population over the season.

### Land use influence on mosquito

Overall, the adjusted pseudo-R^2^ values related to land use were found to be relatively low. In part, a relatively low effect of land use categories could be due to the spatial resolution of land use data being 30m, which may be too coarse to adequately represent variation in mosquito habitat; a finer land use data resolution (∼ 1m) could lead to high adjusted pseudo-R^2^ values [46]. The sample random effect and / or the spatial fixed effects included in the models often showed a relatively high proportion of variance, which could be further evidence that the full impact of land use on mosquito occurrence was not captured in this study. Indeed, our spatial descriptors (db-MEM) likely captured relevant land use structure that were not accounted for in our study such as biogeographic barriers [81].

Wetland appeared to be an important driver of mosquito occurrence or abundance for many species. This is not surprising since wetlands and water constitute a fundamental habitat for the mosquito life cycle, in particular for *An. quadrimaculatus, An. walkeri, Cq. pertubans, Oc. canadensis, Oc. cantator, Oc. japonicus, Oc. trivittatus* and *Ps. ferox* [90, 91], among which are known potential vectors of EEEV and CSV (*Cq. perturbans, Oc. canadensis, Cs. melanura*).

Occurrence of ‘forest’ habitat was also a strong determinant of occurrence and abundance of several mosquito species. Wooded areas generally serve as refuge for adult mosquitoes, especially for the CSV vectors *Oc. excrucians, Oc. communis* and *Oc. stimulans*. These results are consistent with the limited information in the literature which describes, for example, that *Oc. excrucians* can be found in a wide range of habitats according to Wood et al. [90] while Darsie et al [91] mention that these species larvae tend to prefer wooded spring pools and semi-permanent wetlands with abundant vegetation. The ‘forest’ class was negatively associated with *Cx. pipiens-restuans* (presence) and *Anopheles* (abundance), which is consistent with several studies [58, 63, 92]. *Cx. pipiens-restuans* are also known to be urban species although a positive association with ‘urban’ was not detected in our study. Indeed, urban landscapes are very heterogeneous and could themselves be classified into several categories according to the presence of woodlands and aquatic environments. A more detailed characterization of the urban micro-habitat may help better define the relationships between this land use class (urban) and mosquitoes. The negative association between ‘forest’ and the abundance of *An. walkeri* could be explained by the fact that the adults of this species generally stay near their larval aquatic habitats, in contrast to other species of *Anopheles* (*An. quadrimaculatus* and *An. punctipennis*) which leave their breeding sites during the day to seek shelters [91].

The ‘bare ground’ class was negatively associated with mosquito occurrence and abundance, as could be expected as the lack of an aquatic environment and vegetation would prevent completion of the mosquito life cycle.

Intensive agricultural practices, as are common in the study region, could negatively influence some species, including *Anopheles* species and one *Culiseta* species. *Culiseta minnesotae* is naturally adapted to specific marshy environments including sedge and cattail marshes which may be found at the edge of some agricultural fields [91, 93]. However, cattails are considered as invasive plants and are subject to considerable efforts to eliminate them by most farmers, reducing habitat for this species. *Anopheles* species generally prefer pristine freshwater environments such as peat bogs and unpolluted freshwater swamps [90, 91], while wetlands in agricultural land are often contaminated with chemical agents (fertilizers, pesticides, etc.) [94, 95], which may explain the negative association between these species and the agriculture land use class.

The negative relationship between the shrubland class and some mosquito species (*Cq. perturbans, Oc. canadensis, Oc. communis, Oc. intrudens* and *Oc. stimulans*) may be explained by the lack of wetland in this type of habitat, dominated by shrubs, including grasses, herbs, and geophytes.

In summary, we found evidence that mosquito occurrence and abundance are related to land use variables. We detected some common patterns among mosquito species with regard to their response to natural land use variables (water and forest) but more specific response to other land use variables such as agricultural land and shrubland.

## Conclusion

Efforts to connect mosquito distribution and abundance to environmental variables are generally limited to the most regionally relevant species in terms of public health, most being focused on mosquito abundance. However, it is important to determine how multiple species are influenced by these variables since models based on a single or small number of species do not provide a portrait of the whole community including rare disease vectors. This study provides baseline multi-species associations where species respond differently to a given set of environmental conditions. Our study also demonstrates the relevance of including land use in predicting mosquito occurrence and abundance. If temperature and precipitation are primary drivers in mosquito development, appropriate habitat is also required for many species. Our findings suggest that the impact of land use and weather together on mosquito occurrence and abundance is species-specific and may have diverging impacts on whether a species can survive (occurrence) or thrive (abundance). Future research should develop our understanding of mosquito community preferences in terms of habitat and climatic conditions. Several analytical approaches are now available to work at the multi-species level and study the ecological connections between species. One of the main outcome of such study would be to confirm if human-altered environments favor the multiplication of a few species of mosquitoes that are flexible habitat generalists, to while excluding a diversity of more specialized mosquitoes which are becoming increasingly rare. We expect that this imbalance in terms of biodiversity would favor species that are efficient vectors of human disease. Further information about mosquito community ecology could allow vector management strategies to be more targeted with respect to anthropogenic pressures including deforestation, urbanization and agriculture.

## Acknowledgement

We are grateful to all AAFC field teams, Curtis Russel and folks from Public Health Ontario for providing Ontario surveillance data, Julie Légaré for mosquito identification and Christopher Fernandez Prada for lab facilities. We thank the Groupe de Recherche sur l’Épidémiologie des Zoonoses et Santé Publique (GREZOSP) and the Public Health Agency of Canada (PHAC) which funded the study and supported its execution.

## Author Contributions

**Conceptualization:** Miarisoa Rindra Rakotoarinia, Antoinette Ludwig, Patrick Leighton, Nicholas H. Ogden, David Lapen, Guillaume Blanchet, Dominique Gravel.

**Formal Analysis:** Miarisoa Rindra Rakotoarinia, Guillaume Blanchet, Antoinette Ludwig.

**Funding Acquisition:** Antoinette Ludwig, David Lapen.

**Methodology:** Miarisoa Rindra Rakotoarinia, Antoinette Ludwig, Patrick Leighton, Nicholas H. Ogden, David Lapen, Guillaume Blanchet, Dominique Gravel.

**Resources:** Antoinette Ludwig, Patrick Leighton, Nicholas H. Ogden, David Lapen, Guillaume Blanchet, Dominique Gravel.

**Software:** Guillaume Blanchet, Dominique Gravel.

**Supervision:** Antoinette Ludwig, Patrick Leighton, Nicholas H. Ogden, David Lapen, Guillaume Blanchet, Dominique Gravel.

**Visualization:** Miarisoa Rindra Rakotoarinia, Guillaume Blanchet, Antoinette Ludwig.

**Writing – Original Draft Preparation:** Miarisoa Rindra Rakotoarinia, Antoinette Ludwig.

**Writing – Review & Editing:** Miarisoa Rindra Rakotoarinia, Antoinette Ludwig, Patrick Leighton, Nicholas H. Ogden, David Lapen, Guillaume Blanchet, Dominique Gravel.

## Supporting information

S1 File.

(PDF)

## Notes

### Competing Interest Statement

The authors have declared no competing interest.

